# Reactive oxygen species (ROS) triggers unconventional secretion of antioxidants and Acb1

**DOI:** 10.1101/628321

**Authors:** David Cruz-Garcia, Nathalie Brouwers, Vivek Malhotra, Amy J. Curwin

## Abstract

Nutrient deprivation triggers the release of signal-sequence-lacking Acb1 and the antioxidant superoxide dismutase 1 (SOD1). We now report that secreted SOD1 is functionally active and accompanied by export of other antioxidant enzymes such as thioredoxins, (Trx1 and Trx2) and peroxiredoxin Ahp1, in a Grh1 dependent manner. Our data reveal that starvation leads to production of non-toxic levels of reactive oxygen species (ROS). Treatment of cells with N-acetylcysteine (NAC), which sequesters ROS, prevents antioxidants and Acb1 secretion. Starved cells lacking Grh1 are metabolically active, but defective in their ability to regrow upon return to growth conditions. Treatment with NAC restored the Grh1 dependent effect of starvation on cell growth. In sum, starvation triggers ROS production and cells respond by secreting antioxidants and Acb1, which is an important lipogenic signaling molecule in mammalian cells. We suggest that unconventional secretion of antioxidants and Acb1 like activities maintains cells in a form necessary for growth upon their eventual return from starvation to normal conditions.

## Introduction

Unconventional protein secretion (UPS) is defined as the process by which eukaryotic cells export proteins that cannot enter the conventional endoplasmic reticulum (ER) – Golgi complex pathway. This fascinating process was noted upon the cloning of interleukin 1 (IL-1), which lacks a signal sequence for entry into the ER and yet is released by activated macrophages (Auron et al., 1984; Rubartelli et al., 1990). Since then, the repertoire of proteins released unconventionally has expanded considerably. The path taken by this class of proteins to reach the extracellular space varies. Briefly, Type I and II involve direct translocation across the plasma membrane, either via pores or ABC transporters, respectively, while Type III involves incorporation of the cargoes first into an intracellular membrane compartment (Nickel and Rabouille, 2018; Rabouille, 2017). Fibroblast growth factor (FGF)2 and Acyl-CoA binding protein (ACBP/AcbA/Acb1) are two of the best studied examples of unconventionally secreted proteins thus far. FGF2 follows a Type II route, whereas, Acb1 takes a Type III path via that involves a cellular compartment called CUPS, the cytoplasmic proteins Grh1 and a subset of ESCRTs (Cruz-Garcia et al., 2014, 2018; Curwin et al., 2016; Duran et al., 2010; Steringer et al., 2015, 2017). More recently, the export of cytoplasmic enzyme superoxide dismutase 1 (SOD1) was reported to follow the same pathway as Acb1, and both depend on a di-acidic motif (Cruz-Garcia et al., 2017). The release of IL-1ß, however, does not appear to have a unifying theme, and depending on the cell type or stimulus different mechanisms have been proposed that include pore formation via gasdermin D, autophagy-mediated, and via an intermediate membrane compartment (Andrei et al., 2004; Chiritoiu et al., 2019; Dupont et al., 2011; Liu et al., 2016; Rubartelli et al., 1990).

The release of Acb1 and SOD1 is triggered by nutrient starvation upon culturing yeast in potassium acetate. The secreted Acb1 in lower eukaryotes such as yeast and slime mold functions in signaling and regulating the starvation induced cell differentiation program of sporulation (Anjard et al., 1998; Kinseth et al., 2007; Manjithaya et al., 2010). In human brain, ACBP (also known as Diazepam binding inhibitor or DBI) modulates GABA receptor signaling and, more recently in whole animals ACBP has been identified as an important lipogenic signaling factor (Costa and Guidotti, 1991; Gandolfo et al., 2001; Pedro et al., 2019). But what is the physiological significance of secreted SOD1? Here we report that secreted SOD1 is enzymatically active in the extracellular space. Are other antioxidant enzymes secreted along with SOD1? A starvation specific secretome revealed enrichment in cytoplasmic enzymes that function in response to increased reactive oxygen species (ROS) production or oxidative stress. We show that starvation of cells in potassium acetate leads to elevated levels of ROS and this is required for unconventional secretion of the antioxidant enzymes, and Acb1. Therefore, during nutrient starvation in potassium acetate cells sense increased ROS, leading to secretion of active antioxidant enzymes and proteins like Acb1. These secreted activities are necessary to maintain cells in metabolically active form to survive till their return to normal growth conditions.

## Results

### Secreted SOD1 is functionally active

Culturing yeast in potassium acetate promotes the unconventional secretion of SOD1, but is secreted SOD1 functionally active (Cruz-Garcia et al., 2017)? Wild-type and mutant SOD1 linked to the neurodegenerative disorder amyotrophic lateral sclerosis (ALS) are reportedly secreted in a misfolded state when expressed in a motor neuron cell line (Grad et al., 2014), therefore the enzymatic activity of extracellular SOD1 cannot be presumed. We employed a zymography-based assay to directly test whether secreted SOD1 is enzymatically active in yeast cells cultured in potassium acetate. A mild cell wall extraction assay developed to detect starvation specific secretion of Acb1 and SOD1 was performed in non-denaturing conditions (Cruz-Garcia et al., 2017; Curwin et al., 2016). Intracellular and secreted proteins were then separated by native-gel electrophoresis and a standardized in-gel SOD1 activity assay was performed as described previously (Beauchamp and Fridovich, 1971). Our data reveal that both the intracellular and secreted SOD1 are enzymatically active (Figure 1A). A corresponding western blot indicated that secreted pool of SOD1 is as active as its intracellular counterpart, suggesting that most, if not all, secreted SOD1 is functionally active (Figure 1A).

**Figure 1.**
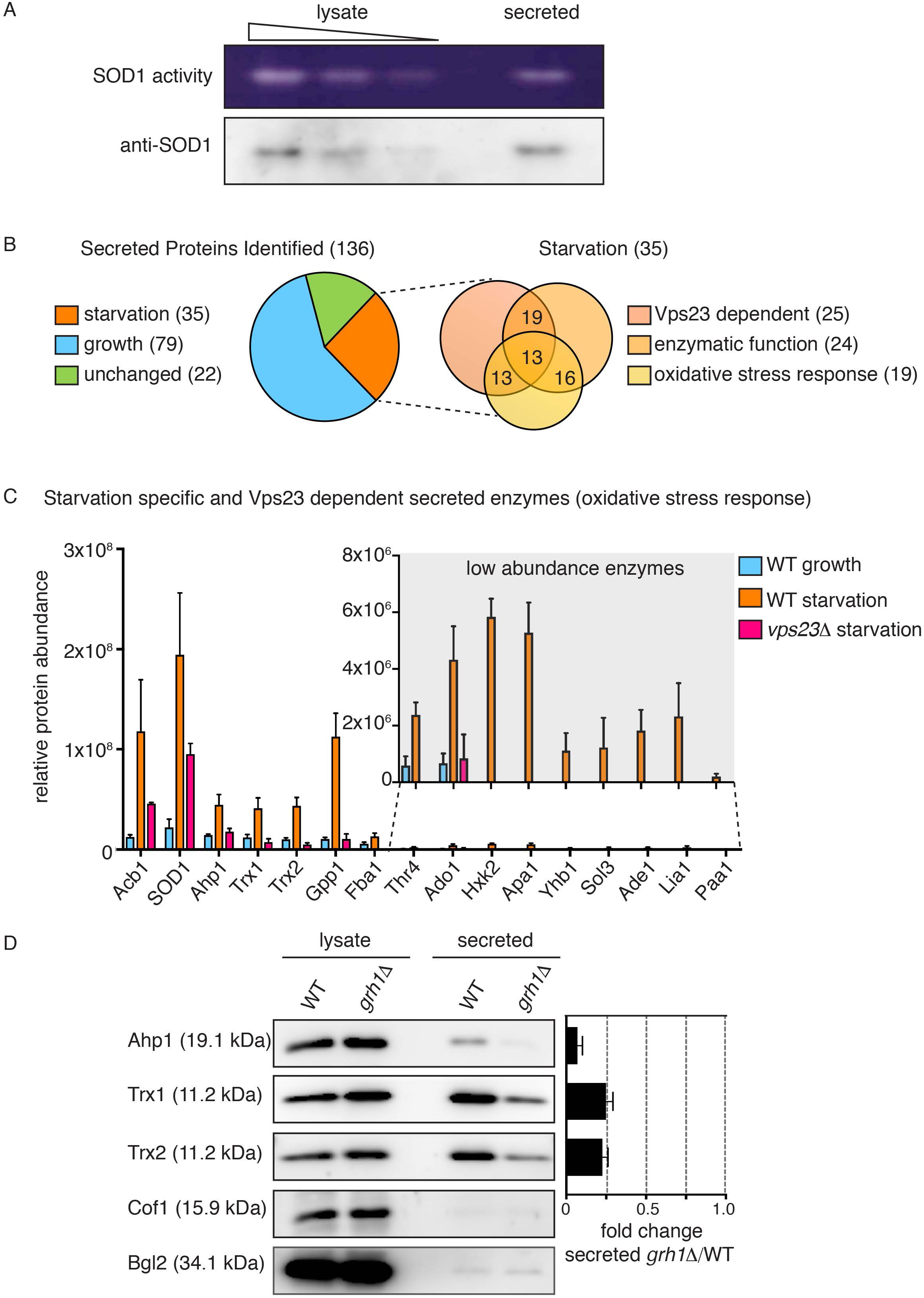
Starvation triggers secretion of cytoplasmic antioxidant enzymes. **(A)** Wild type cells were grown to mid-logarithmic phase, washed twice, and cultured in 2% potassium acetate for 2.5 h. Cell wall proteins (“secreted”) were extracted and concentrated 139x with exchange to TBS (Tris-buffered saline). Separately, a smaller aliquot of the same cells was lysed in non-denaturing conditions in TBS (“lysate”). Lysate and secreted proteins were separated in non-denaturing 12% polyacrylamide gels and either transferred to a nitrocellulose membrane for western blot analysis (“anti-SOD1”) or subjected to the zymography based in-gel SOD activity assay (“SOD1 activity”). Samples of lysate fractions were loaded in decreasing amount, at a dilution of 1/10 to compare with the concentrated secreted fraction. The final relative loading is 80x more in the secreted fraction. **(B)** Mass spectrometry analysis of cell wall extracted proteins (secreted proteins) from wild type cells growing in nutrient-rich conditions (“growth”) versus cultured in potassium acetate for 2.5 h (“starvation”). Cell wall proteins from cells lacking Vps23 were also analyzed under starvation conditions. All three conditions were analyzed in triplicate and statistical analyses were performed to classify the secreted proteins as growth versus starvation specific or unchanged (using a cut-off of a Log2FoldChange of at least -/+ 1, and a *p* value of less than 0.06). Within the starvation specific group, proteins were further classified as dependent on Vps23 or not for their presence in the cell wall, whether they perform an enzymatic activity, and whether they are annotated to affect response to oxidative stress according to the *Saccharomyces* Genome Database (SGD). Complete analyses of proteins identified and the classifications can be found in Tables S1. **(C)** Chart plotting relative protein abundance of the starvation specific/Vps23 dependent enzymes classified as affecting response to oxidative stress. SOD1, Ahp1, Trx1 and Trx2 directly regulate cellular redox balance, while the remaining are key enzymes in various cellular processes known to affect response to oxidative stress such as glycolysis, gluconeogenesis, amino acid or nucleotide biosynthesis, and glycerol metabolism (Acb1 is included only as a reference). The average protein areas from the mass spectrometry analysis (Table S1) of the three triplicates for each protein was calculated, with the corresponding SEM (error bars), for each condition. A zoom of the low abundance enzymes is shown in the inset (note the scale difference). Complete descriptions of the enzymes can be found in Table 1. **(D)** Wild type and *grh1Δ* cells were grown to mid-logarithmic phase, washed twice, and incubated in 2% potassium acetate for 2.5 h. Cell wall proteins were extracted from equal numbers of cells followed by precipitation with TCA (“secreted”). Lysates and secreted proteins were analyzed by western blot and the ratio of the secreted/lysate for the indicated cargo protein was determined and compared to that of wild type in each experiment. Statistical analyses were performed for the indicated unconventional cargo proteins and are represented as fold changes in *grh1Δ* cells compared to wild type (paired student’s t-test). Ahp1=0.07 -/+0.03, *p*=0.0604; Trx1=0.25 -/+0.04, *p*=0.0478; Trx2=0.23 -/+0.03, *p*=0.0065 Error bars indicate SEM, n=3. Cof1 is non-secreted, cytoplasmic protein used to monitor for cell lysis, while Bgl2 is a conventionally secreted cell wall protein. The loading of the secreted fraction is equivalent to 145 x that of the lysate. Based on this the amount secreted compared to the total pool was calculated to be 0.3% for Ahp1, 0.6% for Trx1 and 0.8% for Trx2.

### Identification of a family of secreted antioxidants

Do cells secrete other antioxidant enzymes, like SOD1, upon nutrient starvation? Wild type yeast were cultured in growth or starvation medium, the cell wall proteins were extracted under the conditions described previously (Curwin et al., 2016) and subsequently analysed by mass spectrometry. We also analysed cell wall proteins from cells lacking Vps23, which is required for starvation induced secretion of Acb1 and SOD1, to assist in identification of proteins exported by the same pathway. All analyses were performed in triplicate with statistical evaluation (see Materials and Methods). Using a cut-off of a Log2FoldChange of at least -/+ 1, and a *p* value of less than 0.06, the proteins were classified as starvation versus growth specific, or unchanged (the complete mass spectrometry data and statistical analysis are shown in Table S1). A total of 136 secreted proteins were identified of which 79 were enriched in growth (growth specific), 35 were enriched in starvation (starvation specific), and 22 exhibited no change upon starvation (Figure 1B, Table S1). As expected, the levels of secreted Acb1 and SOD1 were significantly increased upon starvation (log2FC of 3.38 and 2.94 respectively) and their release is Vps23 dependent (Figure 1C, Table S1). Of the 35 starvation specific proteins, 26 required Vps23 for their export (Figure 1B and C, Table S1). Importantly, none of the starvation specific secreted proteins contained a signal peptide for their entry into the ER, as predicted by SignalP 4.1 (Petersen et al., 2011). In fact, only 27 of the 136 total proteins identified contained a signal peptide, all of which were found to be secreted specifically in growth (Table S1).

We were surprised to find that like SOD1, other enzymes directly involved in redox homeostasis were also secreted in a starvation specific manner. These include the thiol specific peroxiredoxin, Ahp1 that reduces preferentially alkyl hydroperoxides, as well as the cytoplasmic thioredoxins of *S. cerevisiae*, Trx1 and Trx2, which function to reduce redox active proteins, including Ahp1 (Holmgren, 1989; Lee et al., 1999). It is also noteworthy that thioredoxin has previously been shown to be secreted unconventionally by mammalian cells (Rubartelli et al., 1992). Closer examination of the starvation specific proteins revealed that 24 are enzymes that function in conserved processes known to affect cellular redox balance (e.g. glycolysis/ gluconeogenesis, amino acid/ nucleotide biosynthesis, and glycerol metabolism) and 19 are actually annotated to function in response to oxidative stress according to the *Saccharomyces* Genome Database (Figure 1B and C, Table 1 and S2). The 15 enzymes involved in response to oxidative stress and dependent on Vps23 for their secretion are highlighted in Figure 1C and Table 1. Therefore, nutrient starvation upon culture of cells in potassium acetate leads to unconventional secretion of a number of enzymes that function, either directly or indirectly, in response to oxidative stress.

**Table 1.**
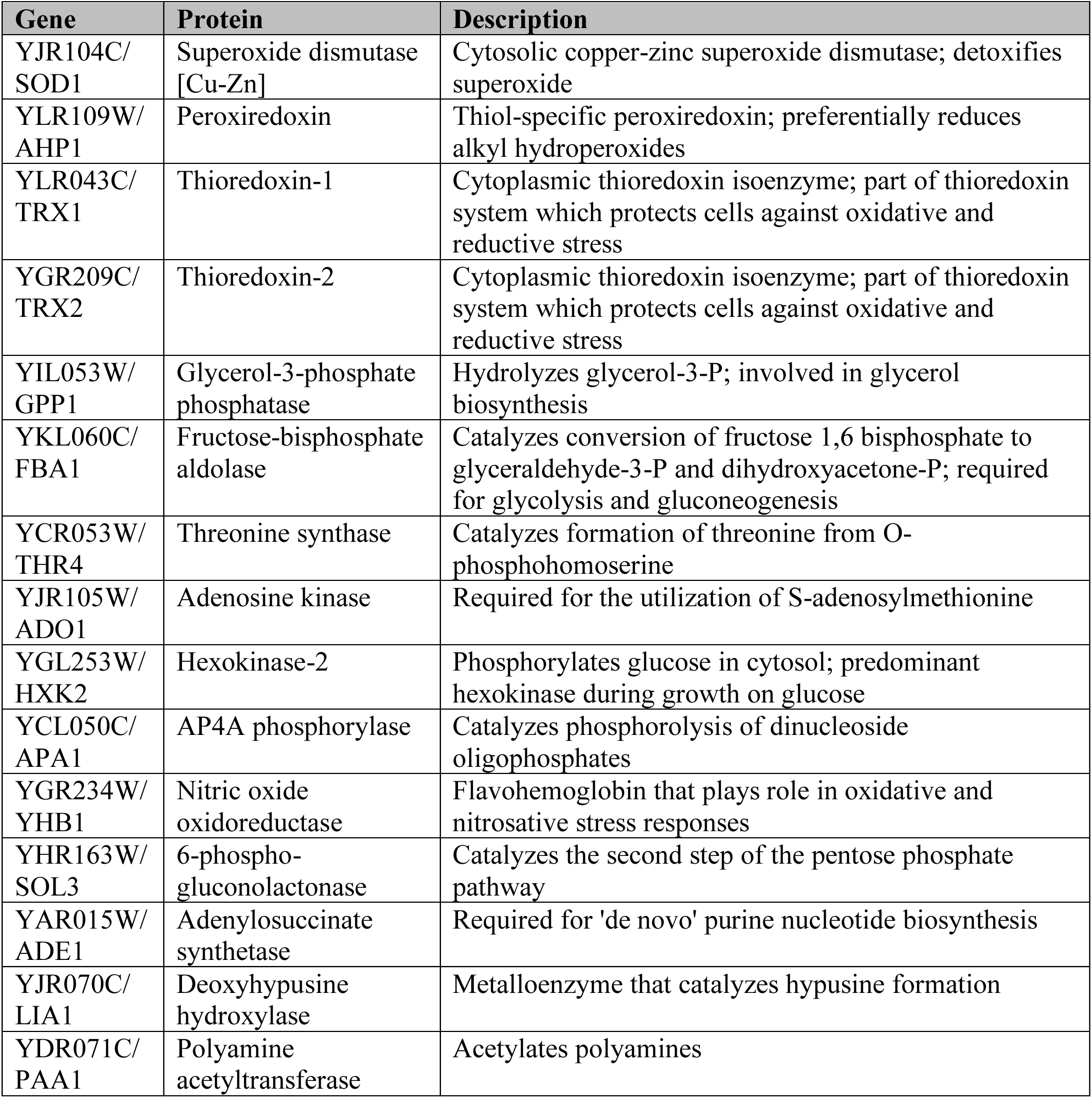
Starvation specific/Vps23 dependent secreted enzymes with direct or indirect functions in response to oxidative stress.

We focussed on three enzymes mentioned above: Ahp1, Trx1 and Trx2. Like SOD1, they function directly in regulating redox homeostasis and are amongst the more abundant enzymes identified in the secretome (Figure 1C, Table S1). Secretion of SOD1 (and Acb1) requires Grh1 (Cruz-Garcia et al., 2017; Curwin et al., 2016), so we first tested the involvement of Grh1 in secretion of these enzymes. A secretion assay of wild-type and *grh1Δ* starved cells revealed that the release of Ahp1, Trx1 and Trx2 upon starvation is also Grh1 dependent (Figure 1D). This, along with the finding that they require Vps23 for their release upon nutrient starvation lends us to conclude that these antioxidant enzymes follow the same pathway of unconventional secretion as SOD1 and Acb1. Secretion of Acb1 and SOD1 was reported to be at ∼1% or less of its total cellular level (Cruz-Garcia et al., 2017; Curwin et al., 2016). Similarly, secretion of Ahp1, Trx1 and Trx2 was a minor fraction of the total protein pool, at 0.3%, 0.6% and 0.8% respectively.

### Starvation leads to ROS production

Based on the known antioxidant function of these enzymes, we hypothesized that starvation increases ROS production, which signals their unconventional secretion. Direct detection of ROS in living cells by fluorescent dyes or expression of intracellular probes is ambiguous (Belousov et al., 2006; Chen et al., 2010; Gomes et al., 2005). Therefore, to confirm the production of ROS during starvation we decided to visualize the location of Yap1, the major transcription factor that responds to multiple forms of oxidative stress by its translocation to the nucleus (Kuge et al., 1997; Schnell et al., 1992). As a control, we exposed yeast cells to 1 mM H_2_O_2_, a strong oxidative stress to the cell, that leads to rapid and complete translocation of Yap1-GFP to the nucleus (Figure 2A and (Delaunay et al., 2000)). Exposure of cells to lower amounts of H_2_O_2_ caused Yap1-GFP translocation, but to a much lesser extent (Figure 2A). In cells cultured in potassium acetate, Yap1-GFP could be detected in the nucleus by 2 hours of starvation and in virtually all cells by 3 hours (Figure 2B). Clearly then, cells cultured in starvation medium generate and sense oxidative stress, but not to the extent observed upon strong oxidative stress, such as exposure to 1 mM H_2_O_2_.

**Figure 2.**
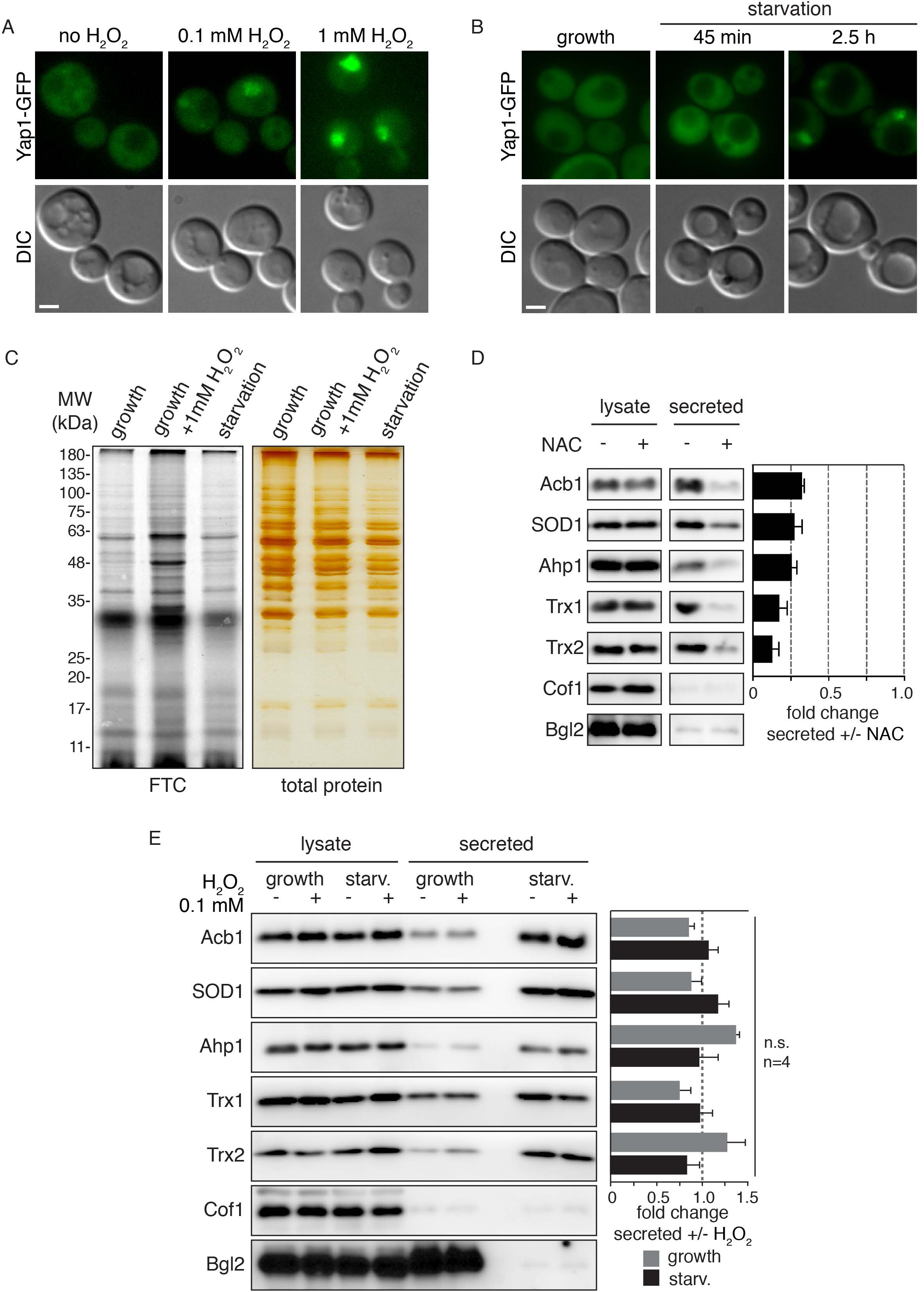
Starvation induces moderate intracellular ROS production, which is required for secretion of antioxidant enzymes and Acb1. **(A)** Yap1-GFP expressing cells were grown to mid-logarithmic phase and incubated in the presence or absence of the indicated amount of H_2_O_2_ for 20 min before visualization by epifluorescence microscopy. Scale bar 2 micron. **(B)** Yap1-GFP expressing cells were grown to mid-logarithmic phase (“growth”), washed twice, and cultured in 2% potassium acetate for 2.5 h (“starvation”). Yap1-GFP was visualized by epifluorescence microscopy. Scale bar 2 micron. **(C)** For detection of total protein carbonylation wild type cells were grown to mid-logarithmic phase (“growth”), washed twice, and cultured in 2% potassium acetate for 2.5 h (“starvation”). Growing cells were also subjected to 1 mM H_2_O_2_ treatment for 1 h as a positive control. Subsequently, cells were lysed, treated with streptomycin sulphate to remove nucleic acids and protein carbonyl groups were labelled with fluorescein-5-thiosemicarbazide (FTC) for 2 hours in the dark. Proteins were precipitated with TCA, unbound FTC was removed, protein pellets were resuspended and separated by SDS-PAGE. Fluorescence was visualized with an Amersham Biosciences Typhoon scanner, while total protein was visualized separately by silver staining. **(D)** Wild type cells were grown to low-logarithmic phase, and either treated with 10 mM n-acetylcysteine (NAC) for 2 h (+) or not (-). Subsequently, equal cell number were washed twice, and incubated in 2% potassium acetate for 2.5 h. Cell wall proteins were extracted and precipitated with TCA (“secreted”). Lysates and secreted proteins were analyzed by western blot. The loading of the secreted fraction is equivalent to 145x that of the lysate. The ratio of secreted/lysate for each cargo was determined and +NAC was normalized to -NAC in each experiment. Statistical analyses were performed for each cargo and are represented as fold change upon NAC treatment (paired student’s t-test). Acb1=0.32 -/+0.02, *p*=0.0335; SOD1=0.27 -/+0.05, *p*=0.032; Ahp1=0.26 -/+0.03, *p*=0.0004; Trx1=0.17 -/+0.05, *p*=0.0597; Trx2=0.13 -/+0.04, *p*=0.002. Error bars indicate SEM, n=3. Cof1 is non-secreted, cytoplasmic protein used to monitor for cell lysis, while Bgl2 is a conventionally secreted cell wall protein. **(E)** For “growth “samples, wild type cells were grown to mid-logarithmic phase and treated (+) or not (-) with 0.1 mM H_2_O_2_ for 1 hour. For starvation (“starv.”) samples, wild type cells were grown to mid-logarithmic phase, equal cell number were washed twice, and incubated in 2% potassium acetate for 2.5 h in the absence (-) or presence (+) of 0.1 mM H_2_O_2_. Subsequently, cells and samples were processed as in (A) to analyze secreted versus lysate pools of the indicated cargo proteins. The same statistical analyses were performed to compare the fold change upon H_2_O_2_ treatment in the growth or starvation conditions. The observed changes were minor and not found to be statistically significant. Error bars indicate SEM, n=4. Cof1 is non-secreted, cytoplasmic protein used to monitor for cell lysis, while Bgl2 is a conventionally secreted cell wall protein.

Carbonylation of proteins and lipids is an irreversible covalent modification that occurs in response to high levels of ROS and is an indicator of overall cellular damage due to excessive ROS production (Fedorova et al., 2014). Protein carbonylation can be detected by labelling total protein extracts with a fluorescent dye FTC (fluorescein-5-thiosemicarbazide) that specifically reacts with the carbonyl moiety on proteins. Again, we used growing cells exposed to 1 mM H_2_O_2_ as a control and compared the level of protein carbonylation in growing and starved cells. Our data reveal that starvation of cells in potassium acetate for 2.5 hours did not lead to any detectable level of protein carbonylation, while cells treated with 1 mM H_2_O_2_ for 1 hour had clearly increased levels of carbonylated proteins, (Figure 2C). The combined evidence shows that cells starving in potassium acetate produce and respond to low, non-damaging amounts of ROS.

### Elevated intracellular ROS is required for secretion

To test whether ROS production is a pre-requisite for unconventional protein secretion upon starvation, we asked if treatment of cells with the antioxidant n-acetylcysteine (NAC) affects secretion. To ensure that NAC is efficiently taken up by cells and to prevent the possibility that it might affect the ability of cells to sense starvation, growing cells were pre-incubated with 10 mM NAC for 2 hours, washed, and resuspended in potassium acetate without NAC, so the starvation period was unchanged from a typical secretion assay. The cell walls were extracted and analyzed as before, revealing that this pre-treatment with NAC was sufficient to significantly reduce secretion of all the unconventionally secreted cargoes tested (Figure 2D). We can thus conclude that starvation of yeast in potassium acetate leads to ROS production, which is required for unconventional secretion of antioxidant proteins. But is elevated ROS alone sufficient to induce secretion? To test this, we asked whether addition of exogenous H_2_O_2_ to growing cells would lead to secretion of the antioxidant enzymes. H_2_O_2_ is the only ROS that can freely cross membranes, while charged ROS, such as the superoxide anion (O_2_^-^) or hydroxyl groups (OH^-^) cannot. We tested addition of 0.1 mM H_2_O_2_, which was sufficient to induce Yap1-GFP translocation to the nucleus to a similar extent as observed upon starvation (Figure 2A and B). Growing and starving cells were exposed to 0.1 mM H_2_O_2_ for 1 hour or the 2.5-hour duration of starvation, and secreted antioxidant enzymes were analyzed by western blot. The results reveal that exogenous addition of 0.1 mM H_2_O_2_ does not induce secretion during growth, nor enhance secretion upon starvation. Therefore, simply creating a pool of cytoplasmic ROS, specifically H_2_O_2_, is not sufficient to induce unconventional secretion of these antioxidants. This indicates the significance of specific species of reactive oxygen, and/or the source of the ROS, in antioxidant secretion.

The major source of ROS in most cell types is the mitochondria, as by-products of aerobic respiration, therefore we decided to investigate mitochondrial organization and function during starvation (Costa and Moradas-Ferreira, 2001; Murphy, 2009; Starkov, 2008). *S. cerevisiae* cells growing in glucose actively repress mitochondrial respiration and rely entirely on glycolysis for energy (known as fermentation) (Ahmadzadeh et al., 1996; Entian and Barnett, 1992). When the carbon source is changed to a non-fermentable one, such as glycerol, ethanol or acetate, cells switch to mitochondrial respiration as the major source of ATP (Boy-Marcotte et al., 1998). Our nutrient starvation conditions consist of culturing cells in potassium acetate, therefore, we expect an acute decrease in ATP production due to lack of glucose, followed by a switch to mitochondrial respiration for ATP synthesis. To monitor the activity of the mitochondria in starvation conditions, we incubated cells with Mito-Tracker, which preferentially labels actively respiring mitochondria (Kholmukhamedov et al., 2013). We also expressed, exogenously, a mitochondrial targeted DsRed fusion protein that labels mitochondria regardless of their activity (Meeusen et al., 2004). In cells grown in glucose, Mito-Tracker and mito-DsRed labelled mitochondria are identified as long tubules (Figure 2A and B). Upon switch to potassium acetate, an increase in mitochondrial membrane potential was immediately apparent, as indicated by greatly increased intensity of Mito-Tracker labelling (Figure 3A). We also noted that mitochondrial morphology was altered, appearing much larger and rounded, as indicated by both the Mito-Tracker dye and mito-DsRed fusion protein (Figure 3A and B). By 2.5 hours of starvation, all cells exhibited mitochondria with increased membrane potential and altered morphology. The increase of membrane potential indicates the source of ROS in starvation is very likely to be the mitochondria. The altered mitochondrial morphology is reminiscent of the phenotype observed in ERMES (ER-mitochondria encounter structure) mutants, where contact with the ER is lost and mitochondrial division is affected (Kornmann et al., 2009; Murley et al., 2013). This begs the question of how functional these mitochondria are, regardless of ROS production. As mitochondria represent the sole source of ATP in these culture conditions, we tested the effect of the chemical 2,4-DNP, that abolishes the proton gradient necessary for the mitochondrial ATP synthase function without directly affecting electron transport (Loomis and Lipmann, 1948). Treatment with 2,4-DNP strongly affected secretion of all cargo proteins tested in a dose dependent manner (Figure 3C). Therefore, the process of unconventional secretion is ATP dependent, but this result also indicates mitochondria are the source ATP in these conditions. Uncoupling the events of ATP and ROS production in the mitochondria is therefore not feasible, as inhibitors will lead to loss of ATP production and some, such as Antimycin A, that inhibits complex III, will exacerbate ROS generation due to accumulation of upstream electron transport chain components (González-Flecha and Demple, 1995).

**Figure 3.**
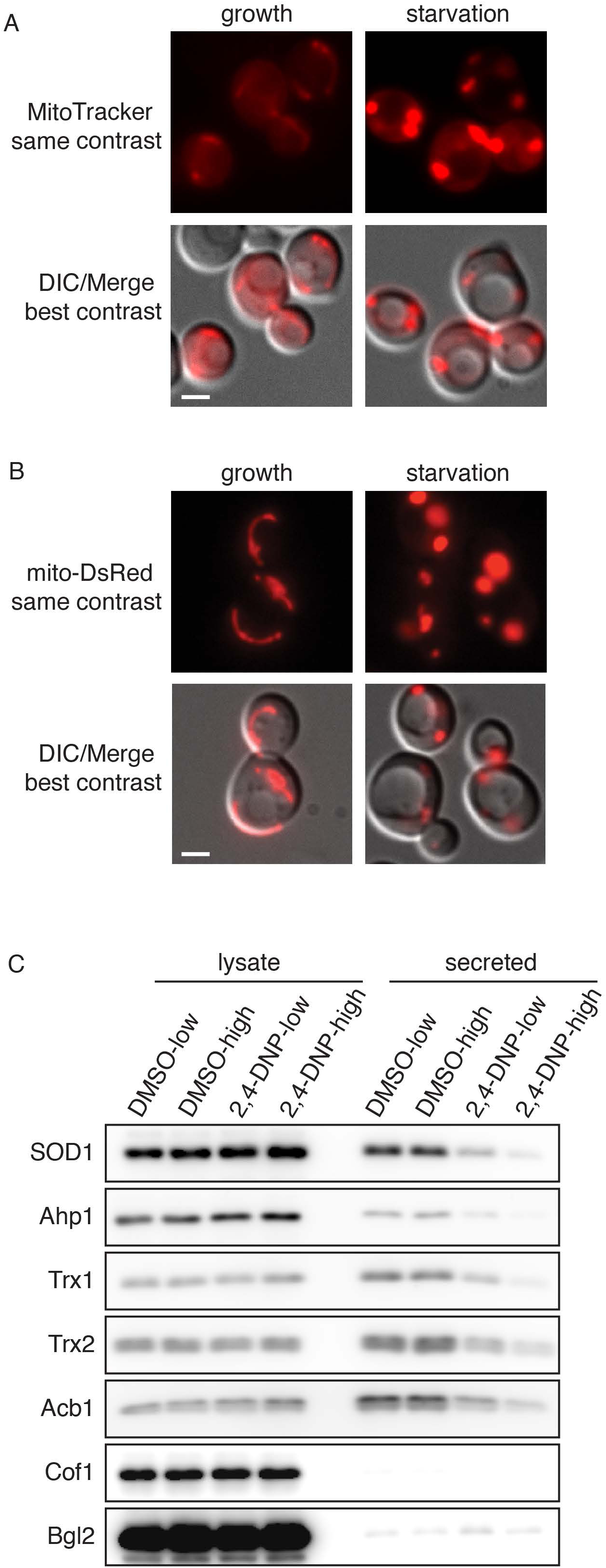
Starvation leads to large, rounded mitochondria with increased activity. **(A)** Wild type cells were grown to mid-logarithmic phase (“growth”), washed twice, and cultured in 2% potassium acetate for 2.5 h (“starvation”). Cells from each condition were labelled with MitoTracker to examine mitochondrial activity and morphology. Cells were visualized by epifluorescence microscopy. Scale bar 2 micron. **(B)** Wild type cells expressing a mitochondrial targeted DsRed construct were cultured and visualized as in (A). Scale bar 2 micron. **(C)** Wild type cells were grown to mid-logarithmic phase, washed twice, and cultured in 2% potassium acetate for 2.5 h in the presence of low or high 2,4-DNP (100 nM and 1 mM, respectively) and the corresponding low or high amount of DMSO carrier as control (1:2000 and 1:500, respectively). Cell wall proteins were extracted from equal numbers cells followed by precipitation with TCA. Lysates and cell wall-extracted proteins were analyzed by western blot. The loading of the secreted fraction is equivalent to 145x that of the lysate. 2,4-DNP treatment strongly inhibited secretion of all proteins analyzed, with low dose ranging from 15-30% reduction, while high dose 2,4-DNP virtually blocked secretion of the cargo proteins, ranging from 3-9% reduction (n=4). Cof1 is non-secreted, cytoplasmic protein used to monitor for cell lysis, while Bgl2 is a conventionally secreted cell wall protein.

Clearly then starvation in potassium acetate for 2.5 hours induces a mild oxidative stress. Western blot of the intracellular pool of antioxidant enzymes indicated there is no upregulation at the unconventionally secreted cargoes within this time frame, implying the cells have not responded to, or sensed yet, the oxidative stress (Figure 2E). This is in line with the fact that Yap1 translocation had just reached ∼100% of the cells by 2.5 hours of starvation (Figure 2B). To get a more comprehensive understanding of how cells respond to starvation, with respect to the intracellular levels of the proteins, we took a quantitative, unbiased approach and performed a total proteome comparison of growing versus starved cells by SILAC (stable isotope labelling with amino acids in cell culture) followed by mass spectrometry. The full data are represented in Table S2 and a volcano plot (Figure S1). None of newly identified secreted proteins were found to change significantly in abundance upon starvation. In fact, the data indicate few dramatic changes in the proteome of cells cultured for 2.5 hours in potassium acetate compared to growing cells; levels of 2 proteins increased, and 9 proteins decreased more than 2-fold of the 990 proteins identified, again indicating cells are still adapting to the new conditions. The 6 largest decreases after starvation were in cell wall modifying enzymes that are secreted conventionally (Figure S1 and Table S2). This is similar to the secretome data (Figure 1 and Table S1) where we observed the absence of signal peptide containing proteins upon starvation. This further shows that during starvation, conventional secretion is inhibited, but there is a net increase in secretion of signal sequence lacking proteins.

### Secretion of antioxidant enzymes during starvation, as well as treatment with NAC, potentiate regrowth upon return to normal conditions

Directly testing the function of the secreted antioxidants is not straightforward. They may provide protection to the cell wall or perhaps, through their activity, could modulate signaling in some way, as ROS can act as a secondary messenger and the secreted antioxidants might be used to control the amplitude of ROS mediated signaling (Dunnill et al.; Hurd et al., 2012). Indeed, the latter hypothesis could explain why such small amounts of enzymes are secreted. Regardless, one would expect some form of advantage to unconventional secretion of antioxidants and proteins like Acb1. Does secretion offer fitness advantage to cells after exposure to starvation? To test this, we measured the ability of cells to regrow after exposure to starvation for 3.5-4 hours. Wild type cells on average had 4.85E+06 CFU/mL, while *grh1Δ* and *vps23Δ* cells produced only 3.25E+05 and 7.17E+05 CFU/mL, respectively (Figure 4A). However, this decrease in CFU/mL is not a result of loss of viability as determined by calcein AM fluorescence. Calcein AM is a membrane permeant dye that only fluoresces in metabolically active or viable cells. Cells were also labelled with propidium iodide (PI) that is non-membrane permeant and only taken up by dead cells. Wild type, *grh1Δ* and *vps23Δ* cells showed 91.8%, 84.5% and 89.9% viable cells, respectively, after 3.5-4 hours of starvation, and only less than 2% dead cells for wild type and *grh1Δ*, while 9.7 % of *vps23Δ* cells were stained by PI (Figure 4B). Therefore, the decrease in CFU/mL observed in *grh1Δ* and *vps23Δ* cells was not due to loss of cell viability. Importantly, we have observed that cells lacking Grh1 can adapt over time, leading to loss of phenotypes linked to unconventional secretion. When newly generated deletion strains were tested immediately in the CFU assay after 3.5 hours of starvation a strong and consistent decrease in CFU/mL was observed in all clones tested. However, the severity of this defect decreased when testing the same clones after longer periods (data not shown). How cells adapt to loss of Grh1is an important new challenge.

**Figure 4.**
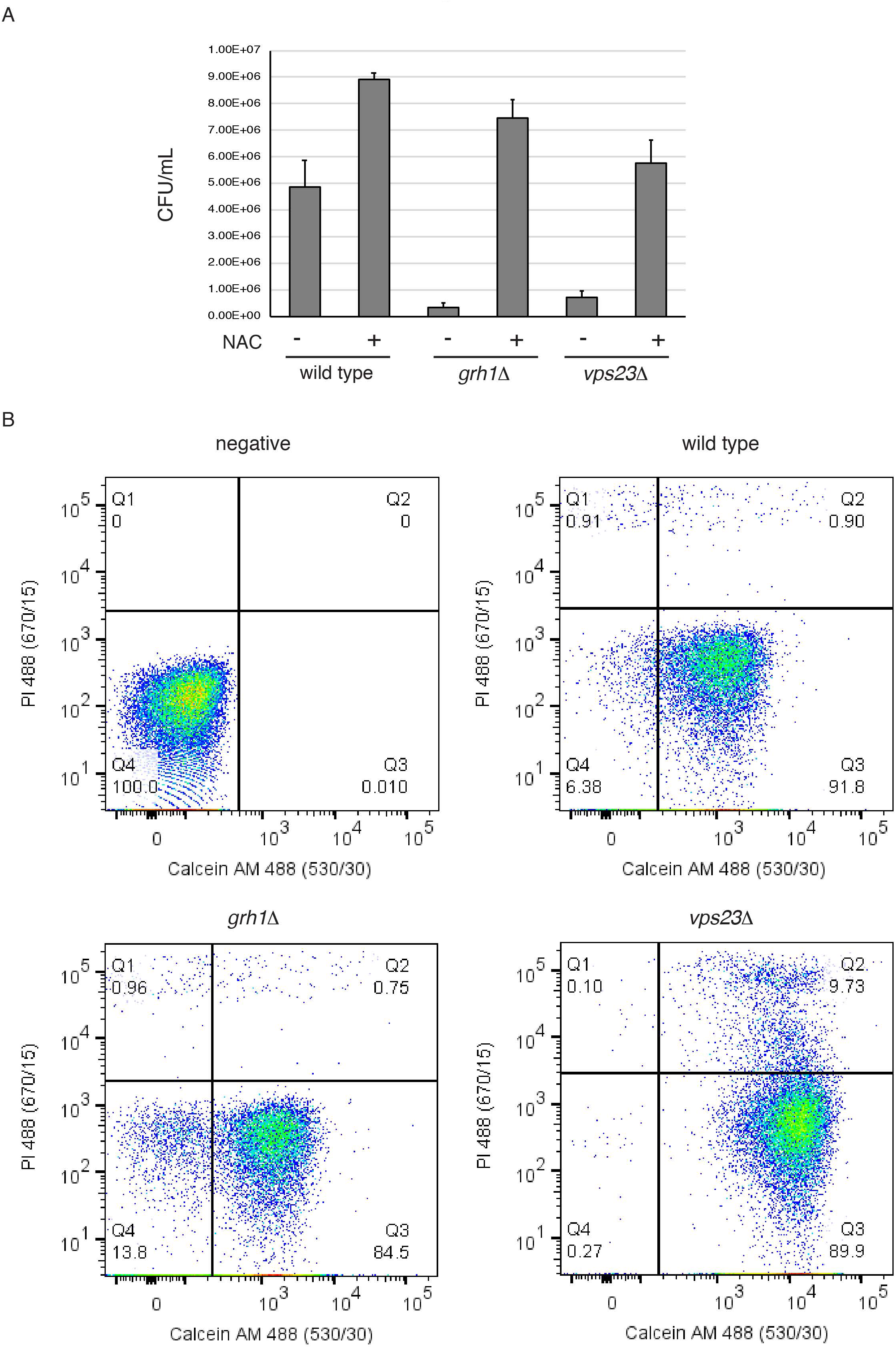
NAC treatment prevents loss of ability to regrow after starvation. **(A)** Wild type, *grh1Δ* and *vps23Δ* cells were grown over night in logarithmic phase, washed twice, and cultured in 2% potassium acetate for 3.5-4 h in the absence or presence of 10 mM NAC. Cell density was normalized to 1 OD/mL during the time of starvation. Subsequently cells were serially diluted to a final dilution of 1:5000 and 100 µL was plated on rich medium. After 2 days of growth at 30°C the number of colony forming units (CFU)/ mL was counted. For wild type cells, n=7, for *vps23Δ* cells, n=3 and for *grh1Δ* cells n=4 of freshly generated deletion strains. All differences were found to be statistically significant by unpaired student’s T-test. Error bars indicate SEM. **(B)** The starving cultures without NAC from (A) were incubated with 5 µg/mL propidium iodide and 1 µg/mL Calcein AM (ThermoFisher Scientific) for 30 min in the dark prior to standard flow cytometry analysis using a FACSCanto (BD Biosciences, San Jose, CA). Data was analyzed with FlowJo software, v10. Representative dot plots are shown, indicating all 3 strains are mostly viable after 3.5-4 h of starvation, based on Calcein AM fluorescence (Y-axis, Q3). Propidium iodide (PI) fluorescence is in the Y-axis.

Interestingly, treatment with NAC increased the CFU/mL of all three strains tested. Wild type cells produced 8.91E+06 CFU/mL (1.8x more), while *grh1Δ* and *vps23Δ* cells now produced 7.45E+06 (18x more) and 5.76E+06 CFU/mL (8x more), respectively, (Figure 4A). The *grh1Δ* cells were more restored to wild type levels than *vps23Δ* cells. This likely reflects the fact that ESCRTs have numerous functions, particularly in starving cells, that may not be specific to UPS, while Grh1 is more specific to this process.

## Conclusion

We have previously shown that a di-acidic motif contained within SOD1 and Acb1 is required for their starvation specific secretion (Cruz-Garcia et al., 2017). In this work, we have focussed on whether the proteins such as SOD1 are functionally active in the extracellular space and the cellular signals that trigger their export. Our findings reveal that SOD1 is secreted in a functionally active form. We have also discovered that cells secrete a number of other enzymes that function together with SOD1 to regulate cellular redox homeostasis. This makes sense because of our findings that culture of yeast in potassium acetate leads to moderate ROS production through increased activity of morphologically changed mitochondria. The cells therefore respond by exporting antioxidant enzymes. Why is Acb1 also secreted in a ROS dependent manner? Acb1 has not been linked to function as an antioxidant. It is however, emerging as an important liopgenic signaling factor secreted upon starvation in mammals (Pedro et al., 2019). We would suggest that starvation triggers ROS production, which then acts as a signal to release a variety of proteins that are composed of antioxidants and proteins that function in other biological pathways that maintain cells in a metabolically active, but otherwise dormant form, till their return to normal growth conditions. ROS and mitochondrial function have been shown to control IL-1ß secretion in mammalian cells (Gabelloni et al., 2013; Jabaut et al., 2013; Zhou et al., 2011). We have shown that Grh1 controls IL-1ß secretion in macrophages via the IRE1 and PERK pathways of the unfolded protein response (UPR) (Chiritoiu et al., 2019). These findings are beginning to reveal the general conserved requirement of Grh1 and ROS in unconventional protein secretion.

Based on our data, we take the opportunity to propose that feeding acetate to yeast causes mitochondria to go into hyper drive, as shown here, which results in production of ROS. As shown recently GRASPs control IL-1ß secretion in macrophages by controlling IRE1/PERK pathway. Cytoplasmic ROS could be sensed as stress signal by GRASPs/Grh1 and conveyed to ER’s UPR pathway, which then navigates cells to respond internally and externally by aptly controlling the production of chaperones and scavengers? How ROS is produced under starvation, the role of mitochondrial form and function, how Grh1 engages in these events, and how ER specific IRE1 and PERK are recruited into the pathway of unconventional secretion are important new challenges to understand the mechanism by which cells cope with starvation till their return to normal growth conditions.

## Materials and Methods

### Yeast strains and media

Yeast cells were grown in synthetic complete (SC) media (0.67% yeast nitrogen base without amino acids, 2% glucose supplemented with amino acid drop-out mix from FORMEDIUM). Wild type strain is BY4741 (*MATa his3Δ1 leu2Δ0 met15Δ0 ura3Δ0*) originally obtained from EUROSCARF. Wild type strain BY4742 was used for the SILAC analysis (*MATalpha his3Δ1 leu2Δ0 lys2Δ0 ura3Δ0*). Deletion strains were from the EUROSCARF *MATa* collection with individual genes replaced by KanMx4. Yap1-GFP was from the Invitrogen collection of C-terminally tagged proteins linked to His3Mx6. Plasmid pVT100U-mtDsRed (J. Nunnari, University of California, Davis) was expressed in strain BY4741 for visualization of mitochondria by microscopy. Multiple new *grh1Δ::KanMx4* strains were generated during this study by amplification of the deletion cassette, with flanking homology, from the EUROSCARF strain and transformation into BY4741.

### Antibodies

All antibodies were raised in rabbit and have been described previously. Anti-Ahp1, anti-Trx1 and anti-Trx2 were the generous gifts of Shusuke Kuge (Tohoku Medical and Pharmaceutical University), Eric Muller (University of Washington) and Yoshiharu Inoue (Research Institute for Food Science, Kyoto University), respectively. Anti-Cof1 was kindly provided by John Cooper (Washington University in St. Louis) and anti-Bgl2 was a gift from Randy Schekman (UC Berkeley). Anti-Acb1 antibody was generated by inoculating rabbits with recombinant, untagged Acb1, purified from bacteria. Specificity of the serum was confirmed by testing lysates prepared from *acb1Δ* cells.

### Cell wall extraction assay

Yeast cells were inoculated at a density of 0.003-0.006 OD_600_/mL in SC medium at 25°C. The following day, when cells had reached OD_600_ of 0.4-0.7 equal numbers of cells (16 OD_600_ units) were harvested, washed twice in sterile water, resuspended in 1.6 mL of 2% potassium acetate and incubated for 2.5 hours. When growing cells were to be analyzed 16 OD_600_ units were directly harvested. The cell wall extraction buffer (100mM Tris-HCl, pH 9.4, 2% sorbitol) was always prepared fresh before use and kept on ice. To ensure no loss of cells and to avoid cell contamination in the extracted buffer, 2mL tubes were siliconized (Sigmacote) prior to collection. Cells were harvested by centrifugation at 3000xg for 3 minutes at 4°C, medium or potassium acetate was removed and 1.6 mL of cold extraction buffer was added. Cells were resuspended gently by inversion and incubated on ice for 10 minutes, after which they were centrifuged as before, 3000xg for 3 minutes at 4°C, and 1.3 mL of extraction buffer was removed to ensure no cell contamination. The remaining buffer was removed and the cells were resuspended in 0.8 mL of cold TE buffer (Tris-HCl, pH 7.5, EDTA) with protease inhibitors (aprotinin, pepstatin, leupeptin (Sigma)) and 10 μL was boiled directly in 90 μL of 2x sample buffer (lysate). For western blotting analysis, 30 μg of BSA (bovine serum albumin (Sigma)) carrier protein and 0.2 mL of 100% Trichloroacetic acid (Sigma) was added to the extracted protein fraction. Proteins were precipitated on ice for 1 hour, centrifuged 16,000xg for 30 minutes and boiled in 50 μL 2x sample buffer. For detection, proteins (10 μL each of lysate or wall fractions) were separated in a 12% polyacrylamide gel before transfer to nitrocellulose (GE Healthcare. For preparation of cell wall extracts for mass spec. analysis, no BSA carrier protein was added and the proteins were precipitated with acetone and not TCA.

### “In-gel” SOD activity assay

After starvation in 2% potassium acetate for 2.5 hours, 8 × 16 OD_600nm_ of cells were subjected to the cell wall extraction protocol as described above. Then, cell wall extracts were pooled and concentrated using Amicon Ultra centrifugal filter 3K devices with buffer exchange into TBS (50 mM Tris-HCl, pH 7.5, 150 mM NaCl) up to a final volume of 75 µl. Also, after incubation in 2% potassium acetate, 16 OD_600nm_ of cells were lysed by bead beating with glass beads in 800 µL of TBS containing protease inhibitors. Lysates were clarified by centrifugation at 16,000 g for 15 min. “In-gel” SOD activity assay was carried out on 12% polyacrylamide gels as described earlier (Beauchamp and Fridovich, 1971). Lysates and concentrated cell wall extracts were separated under native conditions for 7 hours at 40mA. Following completion of the run, the gels were incubated in 2.45 mM Nitroblue Tetrazolium (Sigma) in distilled water for 20 min at room temperature in the dark, followed by additional 15 min incubation with 28 µM riboflavin (Sigma) and 28 mM TEMED (Sigma) in 50 mM potassium phosphate buffer, pH 7.8. Then, the gels were rinsed twice in distilled water, illuminated under a bright white light for 20-30 min and photographed using an Amersham Imager 600. Image processing was performed with ImageJ 1.45r software.

### Preparation of protein samples for secretome mass spectrometry

Samples were reduced with dithiothreitol (27 nmols, 1 h, 37°C) and alkylated in the dark with iodoacetamide (54 nmol, 30 min, 25 °C). The resulting protein extract was first diluted 1/3 with 200 mM NH4HCO3 and digested with 0.9 µg LysC (Wako, cat # 129-02541) overnight at 37°C and then diluted 1/2 and digested with 0.9 µg of trypsin (Promega, cat # V5113) for eight hours at 37°C. Finally, the peptide mix was acidified with formic acid and desalted with a MicroSpin C18 column (The Nest Group, Inc) prior to LC-MS/MS analysis.

### Preparation of samples for total proteome SILAC analysis

Yeast strain BY4742, which is an auxotroph for lysine, was grown for 2 days in 5 ml of synthetic complete media containing either 1 mM L-lysine (light) or 1 mM L-lysine [^13^C_6_^15^N_2_] (heavy) until stationary phase. Cultures were then diluted to OD600 ∼0.005 in 50 ml fresh medium of the same composition until reaching a density of 0.2-0.6. The heavy labelled cells were then washed twice in water and starved in 2% potassium acetate for 2.5 hours. It was not possible to mix the cultures prior to lysis, as the starved cells would sense the nutrients from the growing culture. Therefore, 40 OD units each of growing and starved cells were first pelleted, resuspended in ice-cold lysis buffer (6 M urea, 50 mM Tris–Cl pH 8.0, 0.5% SDS, 0.5% NP-40, 10 mM DDT), and then mixed 1:1. Total cell extracts were prepared in triplicates by glass-bead disruption in lysis buffer. After removal of cell debris by centrifugation at 2000×*g*, proteins were precipitated by the addition of TCA to a final concentration of 20%. The protein pellet was washed with ice-cold acetone which was removed by drying the pellet at 50°C. Proteins were resuspended in a small amount of buffer (6 M urea, 50 mM Tris–Cl pH 7.5) and quantified by the BCA quantification method (Bio-Rad, Hercules, CA). Urea concentration was reduced to 4 M and the pH was adjusted to 8.8 with 25 mM Tris–HCl. Samples were digested with Lys-C in a 1:10 enzyme–protein ratio and desalted using an Oasis Plus HLB cartridge (Waters, Milford, MA).

### Mass spectrometric analysis

The peptide mixes were analyzed using a LTQ-Orbitrap XL (secretome) and an Orbitrap Fusion Lumos (SILAC) coupled to an EasyLC liquid chromatography system (Thermo Fisher Scientific). Peptides were loaded directly onto the analytical column at a flow rate of 1.5-2 µl/min using a wash-volume of 4 times the injection volume, and were separated by reversed-phase chromatography using 25 and 50-cm C18 columns.

The mass spectrometer was operated in data-dependent mode with one full MS scan per cycle followed by the sequential fragmentation of the ten most intense ions with multiple charged.

The mass spectrometry proteomics data have been deposited to the ProteomeXchange Consortium via the PRIDE (Vizcaíno et al., 2016) partner repository with the dataset identifiers PXD010849 (secretome) and PXD016815 (SILAC).

### Data Analysis

Proteome Discoverer software suite (v1.4, Thermo Fisher Scientific) and the Mascot search engine (v2.5, Matrix Science (Perkins et al., 1999)) were used for the secretome peptide identification, whereas SILAC data was analysed with MaxQuant software suite (v1.5.4.3) and the Andromeda search engine. Samples were searched against a *S. cerevisiae* database (version February 2014) containing a list of common contaminants and all the corresponding decoy entries. Trypsin (secreome) or Endopeptidase LysC (SILAC) was chosen as enzyme, and carbamidomethylation (C) was set as a fixed modification, whereas oxidation (M), acetylation and (N-terminal) were used as variable modifications. The SILAC samples also included 13C6,15N2-Lys (+8) as variable modification. Searches were performed using a peptide tolerance of 4.5-7 ppm, a product ion tolerance of 0.5 Da and the resulting data files were filtered for FDR < 5 %. Volcano plot of SILAC data was generated using R studio Bioconductor package “EnhancedVolcano” (Blighe et al., 2019).

### Detection of total protein carbonylation

For detection of total protein carbonylation wild type cells were grown to mid-logarithmic phase (“growth”), washed twice, and cultured in 2% potassium acetate for 2.5 h (“starvation”). Growing cells were also subjected to 1mM H_2_O_2_ treatment for 1 h as a positive control. Subsequently cells were lysed by bead beating in carbonylation buffer (20 mM Na-phosphate pH 6, 1 mM EDTA, 6 M Urea, plus protease inhibitors) and cell extracts were cleared by centrifugation at 10,000 x g, for 10 min at 4°C. Supernatants were treated with 1% streptomycin sulphate for 5 min to remove nucleic acids, and cleared again by centrifugation at 10,000 x g for 5 min at 4°C. Extracts were diluted to 1 µg/µL in carbonylation buffer and 200 µL was labelled with 50 mM fluorescein-5-thiosemicarbazide (FTC) (Fluka) for 2 h in the dark. Proteins were precipitated with TCA, unbound FTC was removed by washing 5 times in ice cold ethanol:ethyl acetate (1:1). Protein pellets were resuspended 50 µL dissolving buffer (20 mM Na-phosphate pH 8, 1 mM EDTA, 8 M Urea) and samples were prepared with 5x sample buffer without bromophenol blue for separation in a 12% SDS-acrylamide gel by running 5 µg of protein per lane, in 2 separate gels. Total protein was visualized in one gel by silver staining, while fluorescent carbonylated proteins were visualized in the second gel using an Amersham Biosciences Typhoon scanner.

### Fluorescence Microscopy

After incubation in the appropriate medium, cells were harvested by centrifugation at 3000 x g for 3 min, resuspended in a small volume of the corresponding medium, spotted on a microscope slide, and imaged live with a DMI6000 B microscope (Leica microsystems, Wetzlar, Germany) equipped with a DFC 360FX camera (Leica microsystems) using an HCX Plan Apochromat 100x 1.4 NA objective. Images were acquired using LAS AF software (Leica microsystems) and processing was performed with ImageJ 1.47n software. For MitoTracker staining 10 OD_600nm_ of cells were harvested by centrifugation at 3,000 *g* for 4 min, resuspended at 5 OD_600nm_ /ml in 100 nM MitoTracker Red CMXRos (Molecular Probes) diluted in phosphate-buffered saline, and incubated in the dark for 30 min. Then, ∼1.5 OD_600nm_ of cells were harvested by centrifugation at 3,000 *g* for 3 min, resuspended in a small volume, spotted on a microscopy slide, and subsequently live-imaged.

### Online supplemental material

Table S1 shows the complete secretome mass spectrometry data, including subsequent statistical analysis, and the classification of each identified protein as “growth specific”, “starvation specific” or “unchanged”; “Vps23 dependent” or “Vps23 independent”; or as containing or lacking a signal peptide. Table S2 shows the complete data of total proteome comparison of growth versus starvation by SILAC and subsequent statistical analysis. The 35 starvation specific secreted cargoes have also been highlighted separately, indicating no significant changes. Figure S1 shows a volcano plot of the SILAC data, highlighting minor changes in the proteome after starvation.

## Supporting information

Supplemental Table 1 (secretome)

Supplemental Table 2 (whole proteome SILAC)

## Acknowledgements

We thank members of the Malhotra lab for their valuable discussions. We also thank Dr. Elena Hidalgo (UPF) and her lab members for their expertise and technical assistance related to oxidative stress. V. Malhotra is an Institució Catalana de Recerca i Estudis Avançats professor at the Centre for Genomic Regulation. This work was funded by grants from the Spanish Ministry of Economy and Competitiveness (BFU2013-44188-P and BFU2016_75372-P to VM). We acknowledge support of the Spanish Ministry of Economy and Competitiveness, through the Programmes “Centro de Excelencia Severo Ochoa 2013-2017” (SEV-2012-0208 & SEV-2013-0347). The mass spectrometry data has been acquired at the CRG/UPF Proteomics Unit, which is part of Proteored, PRB3 and is supported by grant PT17/0019, of the PE I+D+i 2013-2016, funded by ISCIII and ERDF. We also acknowledge the CRG/UPF FACS facility. This work reflects only the authors’ views, and the EU Community is not liable for any use that may be made of the information contained therein.

---

**Figure S1.**
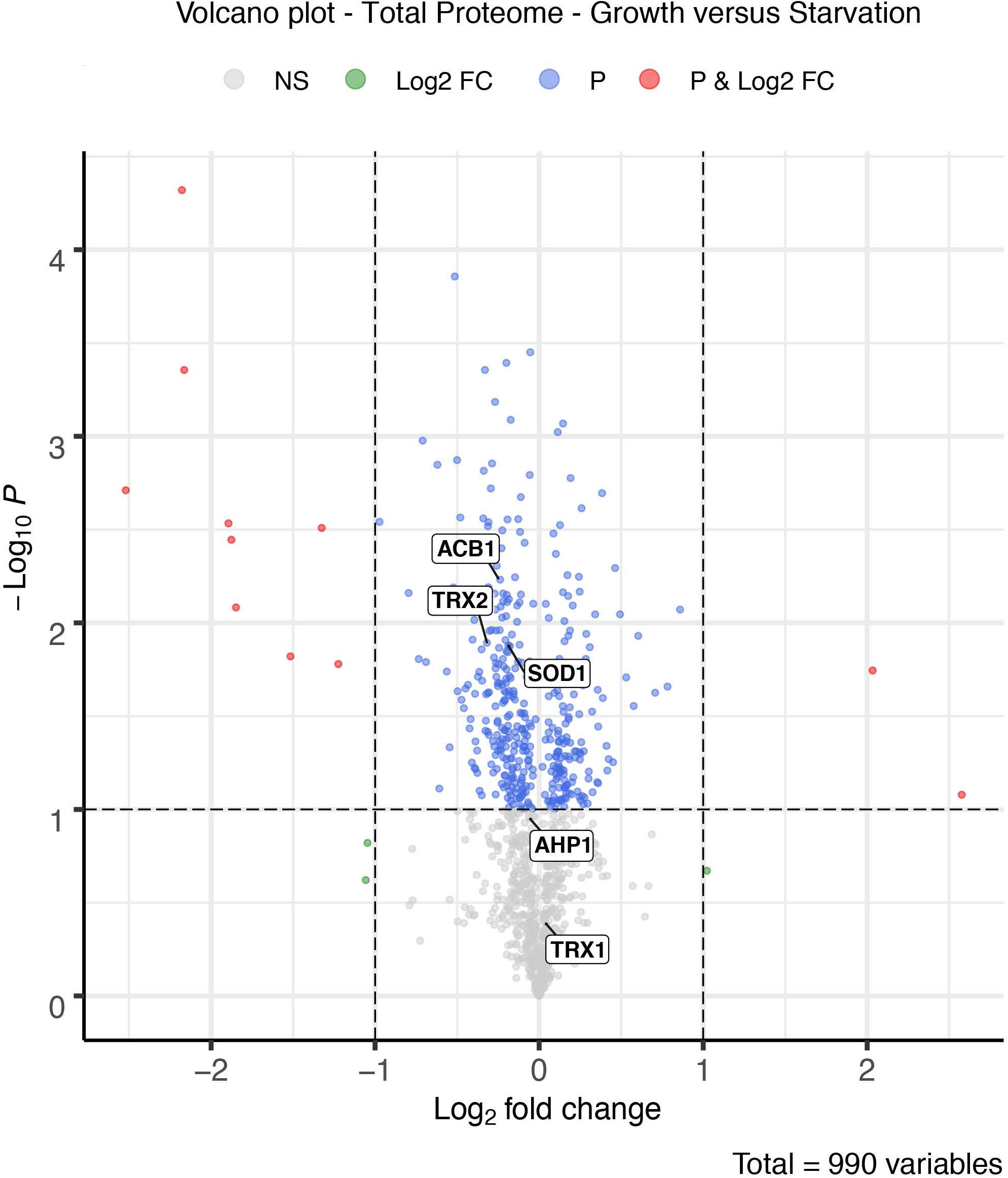
Volcano plot representing the SILAC whole proteome comparison. Positive Log2Fold changes represent increases and negative Log2Fold changes represent decreases after 2.5 h of starvation (n =3). Complete details in materials and methods.

